# DeepLPI: a novel deep learning-based model for protein-ligand interaction prediction for drug repurposing

**DOI:** 10.1101/2021.12.01.470868

**Authors:** Bomin Wei, Yue Zhang, Xiang Gong

## Abstract

The substantial cost of new drug research and development has consistently posed a huge burden for both pharmaceutical companies and patients. In order to lower the expenditure and development failure rate, repurposing existing and approved drugs by identifying interactions between drug molecules and target proteins based on computational methods have gained growing attention. Here, we propose the DeepLPI, a novel deep learning-based model that mainly consists of ResNet-based 1-dimensional convolutional neural network (1D CNN) and bi-directional long short term memory network (biLSTM), to establish an end-to-end framework for protein-ligand interaction prediction. We first encode the raw drug molecular sequences and target protein sequences into dense vector representations, which go through two ResNet-based 1D CNN modules to derive features, respectively. The extracted feature vectors are concatenated and further fed into the biLSTM network, followed by the MLP module to finally predict protein-ligand interaction. We downloaded the well-known BindingDB and Davis dataset for training and testing our DeepLPI model. We also applied DeepLPI on a COVID-19 dataset for externally evaluating the prediction ability of DeepLPI. To benchmark our model, we compared our DeepLPI with the state-of-the-art methods of DeepCDA and DeepDTA, and observed that our DeepLPI outperformed these methods, suggesting the high accuracy of the DeepLPI towards protein-ligand interaction prediction. The high prediction performance of DeepLPI on the different datasets displayed its high capability of protein-ligand interaction in generalization, demonstrating that the DeepLPI has the potential to pinpoint new drug-target interactions and to find better destinations for proven drugs.

## Introduction

Introducing a new drug to the market has been characterized to be risky, time-consuming, and costly[1][2]. Drug discovery is the first phase of drug research and development (R&D) that starts with identifying targets of an unmet disease such as proteins, followed by creating and optimizing a promising compound that can interact with the targets efficiently and safely. This step usually involves hundreds and thousands of compounds, yet only about 8% of which as drug leads can enter the phase of the *in vitro* and *in vivo* preclinical research [3]. To shorten the duration and improve the success rate in the phase of drug discovery, drug repurposing has become a hotspot of new drug research and development over the past few years [1][4], which intends to find an effective cure for a disease from a large amount of existing and approved drugs that were developed for other purposes [1]. For example, prednisone was originally developed for the treatment of inflammatory diseases, but it is likely to be effective against Parkinson’s disease as well [5]. In midst of all the drug repurposing methods, *in silico* computational-based methods to screen pharmaceutical compound libraries and identify drug-target interactions (DTIs) or protein-ligand interactions (PLIs) have gained increasing attention and made significant breakthroughs due to the development in high performance computational architectures and advances in machine learning methods.

Over the last decade, a variety of machine learning-based models have been developed to identify PLIs from millions of ligands and proteins. One type of models utilized 3D structures of proteins and drug molecules aiming at capturing interaction details in predictions of the drug-target binding affinity[6], such as Atomnet[7] and SE-OnionNet[8], but insufficient 3D protein structure data limited the practicability, generalizability, and accuracy [9, 10, 11]. To exploit the vastly available protein sequencing data, a new type of model calculates human-selected features and predicts drug-target interactions with conventional machine learning [12, 13]. The disadvantages of these methods are that they not only require much domain knowledge but also possibly lead to a loss of the information about raw protein-ligand interactions due to limited features. Deep learning-based models can automatically learn highly complex and abstract level of features from large-scale datasets without extensive manual creation. Yet the recent development considers only simple encoding of input letter information [14, 15, 16]. Without the contextual information, this type of model may not capture the complex protein features and thus have limited accuracy and generalizability.

Here, we propose DeepLPI, an innovative deep learning-based model to predict proteinligand interaction using the simple formats of raw protein 1D sequences and 1D ligands (i.e., drug molecular) SMILES (**S**implified **M**olecular **I**nput **L**ine **E**ntry **S**ystem) strings as inputs, rather than manual-generated features or complex 3D protein structures. To capture contextual information in the sequence data, we first respectively employ Natural Language Processing-inspired, pre-trained models of Mol2Vec[17] and ProSE[18] to embed drug SMILES strings and protein FASTA sequences as numeric vectors. These embedded numeric vectors are then fed into two blocks, each of them consisting of two modules termed head convolutional module and ResNet-based convolutional neural network (CNN) module, to encode proteins and drug sequences, respectively. The encoded representations are concatenated into a vector and further fed into a bi-directional long short-term memory (biLSTM) layer, followed by three fully connected layers. We download the BindingDB dataset [19] and Davis [20] dataset to train the DeepLPI model and adjusted the hyperparameters and internally independently evaluate its performance towards PLI prediction. We further transformed the model on a COVID-19 3CL Protease [21, 22] dataset for externally assessing the prediction ability of DeepLPI. To benchmark our model, we compared DeepLPI with the start-of-the-art methods of DeepDTA [15] and DeepCDA [16] towards protein-ligand interaction. The prediction performance is quantitively represented in terms of area under the receiver operating characteristic curve (AUROC), sensitivity, specificity, positive (PPV), predictive value, and negative predictive value (NPV). The high performance of our DeepLPI towards protein-ligand interaction prediction suggests that our model has the potential to accurately identify protein-ligand interaction and hence, promote the new drug development.

## Methods

### Dataset and data preprocessing

We use the BindingDB[19] and Davis [20] datasets to train and evaluate our DeepLPI model. We also use the COVID-19 3C-like Protease dataset from Diamond Light Source [21, 22] for further assessment. All datasets are publicly accessible. The BindingDB is a continually updating database that contains 2,278,226 experimentally identified binding affinities between 8,005 target proteins and 986,143 small drug molecules up to July 29, 2021. We first apply the following criteria to compile the dataset for the development of our model (**Fig. S1**): (1) excluding binding interactions with multichain protein complexes because it is not capable of identifying which chain of the protein interacts with the molecular; (2) retaining binding interactions only represented by *K_d_* value and it means that other measurements in the form of *IC*50 or *K_i_* values are removed; (3) keeping common drug molecules and target proteins occurring in at least three and six interactions in the entire dataset [9], respectively; (4) removing data with invalid *K_d_* values and removing duplicated data entries. For example, we notice that some data used “>” and “<” in the labeled values to indicate ranges, so directly exclude them for the subsequent analysis. Additionally, there are some zeros in the values which should not appear based on the definition of binding affinity measurement of *K_d_*. Thus, we treat them as invalid values and simply removed them; As a result, a total of 36,111 interactions with 17,773 drug molecules and 1,915 protein targets are finally used in developing our model. (5) As a binary classification problem in this study, we use label 1 to represent a pair of protein and ligand being active if their corresponding *K_d_* value is less than 100 nM and use label 0 to represent a pair being inactive if their *K_d_* value is greater or equal to 100mN since a greater dissociation constant means weaker binding. In this case, 59.9% of data are labeled active, and 40.1% of data are labeled inactive (**Fig. 1a and 1b**). The median (standard deviation, [minimum, maximum]) lengths of drug molecular SMILES strings and protein sequences are 52 (45.81, [1, 760]) and 445 (456.1, [9, 7096]) (**Fig. 1c and 1d**).

**Figure 1.**
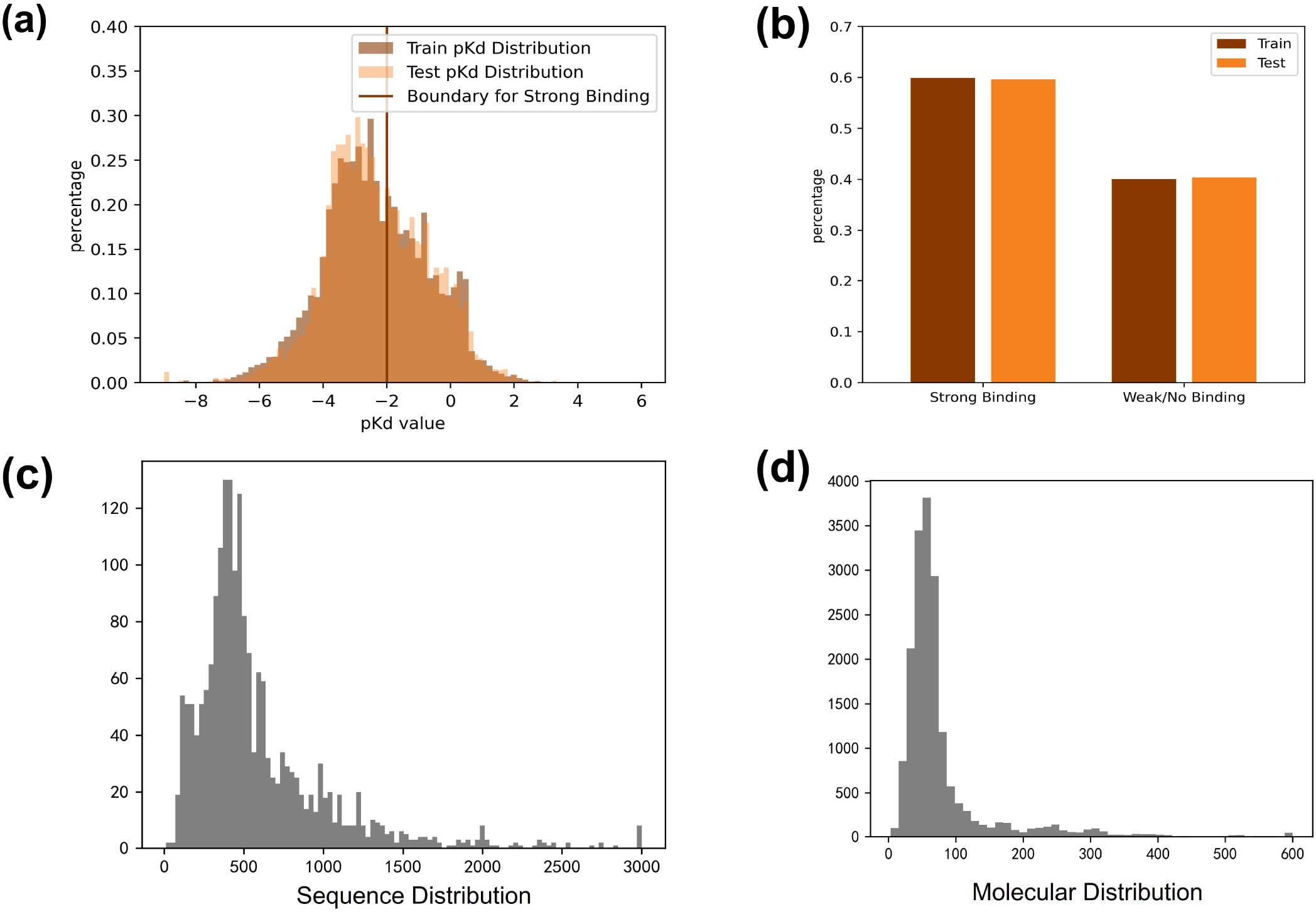
Distribution of BindingDB data used to develop the DeepLPI model. (a) Distribution of the *pK_d_* values and the threshold for determining active/inactive. (b) distribution interaction in binary classes (c) Distribution of lengths of drug molecular SMILES strings. (d) Distribution of lengths of protein sequences.

We then randomly select 85% of the pre-processed BindingDB dataset as the training set and the remaining 15% as the internal independent testing set to train and evaluate our DeepLPI model. To optimize hyperparameters, we further allocate 10% of all data from the training set for validation during the training phase, and the rest are used as a training subset (i.e., 75% of all data). The AUROC and four classification metrics (sensitivity, specificity, PPV and NPV) are computed.

To characterize the model generalizability, different drugs and proteins were selected and reserved for the testing set (**Fig. S2**). “Drug unseen” testing set consists of drugs not seen in the training set, “Protein unseen” testing set consists of proteins not seen in the training set, and “None seen” testing set consists of drugs and proteins neither seen in the training set.

The reason for choosing *K_d_* rather than other binding measurements is for enabling our model performance to be reasonably compared on Davis dataset, which contains interactions of 442 unique proteins and 68 unique compounds. The Davis dataset only reports *K_d_* values of the kinase protein family and the relevant inhibitors. We used the same protocol to obtain the class label as we did above. The Davis dataset was referenced from the Davis work [20] and downloaded from the URL therein. The dataset contained duplicated data entries where the drug-protein pairs are the same, but the binding affinity values are different, potentially due to the experiment conditions. We keep only one entry in each group of duplicates. This doesn’t affect the balance of the dataset because according to our binary threshold, all data entries in the same duplicate group in fact have the same binary label. After the treatment, there are 24,548 interaction data entries. We split them into training, validation, and testing sets according to the same method described above.

To find effective drugs for SARS-CoV-2, we applied our model on a COVID-19 dataset where 879 small molecule drugs were tested on the SARS-COV-2 3C-like protease. The experiment measured EC50 results. For classification, we label 1 to indicate drug-protease active if EC50 is less than 30 nM [23] or 0 representing inactivity. The data is retrieved from a large XChem crystallographic fragment screen against SARS-CoV-2 main protease at high resolution from MIT AiCures. [22] Among those data, 78 are active according to the threshold.

### Model Design

#### Overview of DeepLPI model

The proposed DeepLPI consists of eight modules **(Fig. 2)**, including two embedding modules, two head modules, two ResNet-based CNN modules, one bi-directional LSTM (biLSTM) module, and one multilayer perceptron module (MLP). DeepLPI employs raw molecular SMILES strings and protein sequences as inputs, which are first represented as numeric vectors using the pretrained models of Mol2Vec and ProSE, respectively. The embedded vectors for the drug SMILES and the protein sequences are then fed into the respective head module and ResNet-based CNN module to extract features. The feature vectors for the inputs of drug molecules and protein targets are concatenated, pooled (maxpooling operation), and then encoded by a biLSTM layer. Subsequently, the encoded vectors are finally fed into an MLP module, and the final output is activated through a sigmoid function for binary classification to predict active/inactive labels.

**Figure 2.**
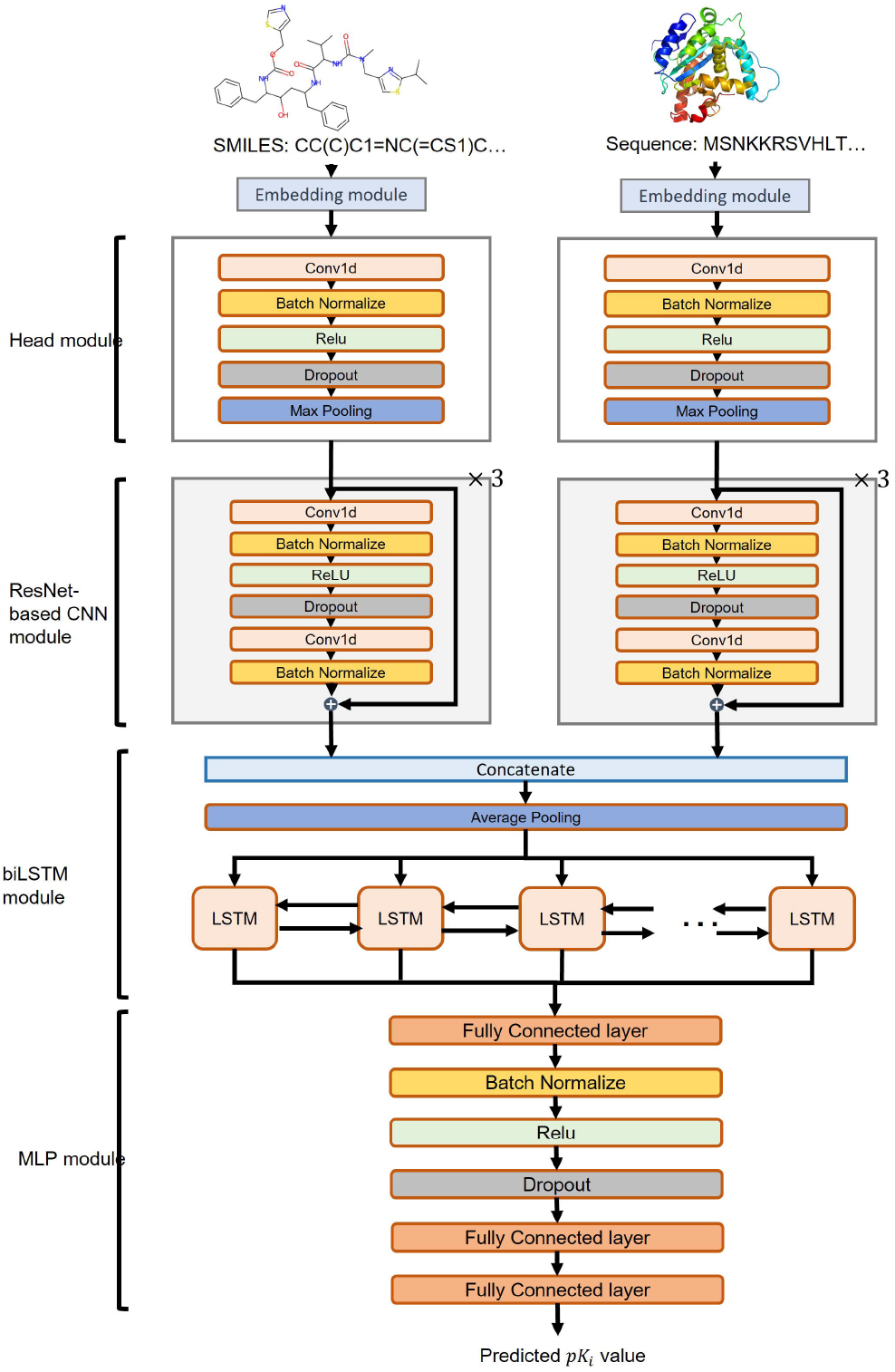
The overview of the DeepLPI model.

##### Embedding module

To utilize the raw drug molecular SMILES string and protein sequence as inputs to the DeepLPI model, we firstly encode them into numeric vector representations using the pre-trained embedding models Mol2Vec[18] and ProSE[19], respectively. Mol2Vec is an unsupervised deep learning-based approach to convert a molecule into a numeric vector representation. Inspired by natural language processing (NLP) techniques, Mol2Vec regards the molecular substructures obtained by the Morgan identifier [23] as “words” and the compound as “sentences”, and then encodes them into dense vector representations based on a so-called corpus of compounds. On the other hand, the ProSE is a deep learning-based method developed to represent protein sequences into numeric vectors that encode protein structural information. It first translates a protein sequence into a list of specific alphabets (as a “sentence”) which map similar amino acids (as “words”) into close numbers. Then, the ProSE model encodes the words into numeric vectors.

We utilize the pre-trained Mol2Vec (download link: https://github.com/samoturk/mol2vec) and ProSE (download link, https://github.com/tbepler/prose) to obtain vector representations with a fixed length for the drug molecular compound and protein, respectively.

##### Head module and ResNet-based CNN module

After the embedding, we separately feed the drug molecular SMILES string vector and protein sequence vector each into the head modules with the same network architecture. The head module contained the following layers: 1D convolutional, batch normalization, nonlinear transformation (with the rectified linear unit, i.e., ReLU activation), dropout, and max-pooling.

Subsequently, two ResNet-based CNN modules are connected to the corresponding head module to further encode the information of input. Suppose *x* is the input into a ResNet-based block, the output of stacked layers is called residual, denoted as *F*(*x*), we then calculated ResNet-based block output with equation *H*(*x*) = *F*(*x*) + *x* [24]. Similar to the head module, the two ResNet-based CNN modules had the same network architecture. Specifically, each ResNet-based CNN module consists of three consecutive ResNet-based blocks, and each block comprises two branches, where the right branch is known as “shortcut connection”; and the left branch is known as a residual network that contains several stacked layers, including a 1D convolutional layer, a batch normalization layer, a ReLU layer, a dropout layer, another 1D convolutional layer, and one more batch normalization layer in sequence.

##### biLSTM module and MLP module

In the biLSTM module, we first concatenate the outputs of features extracted by the two ResNet-based CNN modules, following with an average-pooling layer. The biLSTM, which stands for **bi**directional **l**ong **s**hort-**t**erm memory, can learn long-term dependency from inputs. This network processes the input twice, once from starting to the end and once the reverse way and thus can balance the molecular information and the protein information. Finally, the outputs on both sides of biLSTM will be combined as the output vector.

In the MLP module, we flatten the output vector of biLSTM and fed it into three stacks of consecutive FC layers. Finally, the output is passed through a sigmoid function for binary classification to predict 1/0 labels.

#### Loss function

We treat the prediction as a classification task, predicting whether the drug and protein has a strong binding or a weak binding. For the *n* pairs of molecular SMILES strings and protein sequences the loss function of the DeepLPI model was given by:

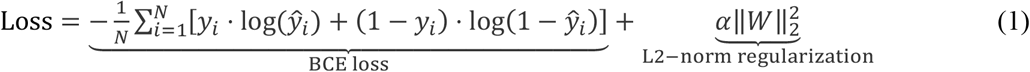

where *y_i_* ∈ {0,1} is class label representing whether or not binding interaction of a input pair of protein and ligand sequences *i*. 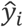 is the probability of interaction prediction for the input pair *i* by our model, 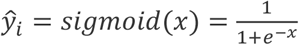, *x* is the output of the MLP module of our model. *W* is the trainable weight matrix in our model. *α* is decay rate and we set it as 0.8 in this study.

#### Evaluation metrics

We calculate five metrics including area under the receiver operating characteristic curve (AUROC), sensitivity, specificity, positive predictive value (PPV), and negative predictive value (NPV) to evaluate the performance of our model.

1. Sensitivity:

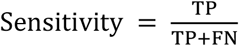

where TP represents true positives, FP represents false positives, TN represents true negatives, FN represents false negatives.
2. Specificity:

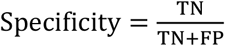
3. Positive predictive value (PPV):

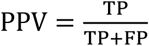
4. Negative predictive value (NPV):

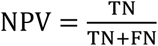

#### Experiment setup

Model training was done in Aliyun Cloud Computing. The node CPU used Intel(R) Xeon(R) Platinum 8163 (2.50GHz). An Nvidia Tesla T4 GPU is supplied. The sources code is available on Github. The model is implemented using the PyTorch library (version 1.8.1). The source code of training and evaluating DeepLPI and the requirements are available on GitHub (https://github.com/David-BominWei/DeepLPI).

#### Parameters setting for training DeepLPI

We use Kaiming Initialization to initialize DeepLPI network weights [25]. The Adam optimizer[26] is also employed with default parameters of *β*_1_ = 0.9 and *β*_2_ = 0.999 as an optimization algorithm to train our model. Furthermore, we use a batch size of 256 and initialize the learning rate at 0.001 with a decay rate of 0.8 for every 10 epochs. The maximum number of epochs is 100 epochs for BindingDB training and 50 for Davis training. All settings for the parameters implemented in our DeepLPI model are demonstrated in **Table S1**. It should be noted that we use the default parameter values for the pre-trained Mol2Vec and ProSE, and we yield vector representations with a fixed length of 300 for the drug molecules, and two lengthes of 100 and 6,165 for the target proteins. The 6,165-element representation for protein were tested to outperform the 100-element representation, and thus in the article we report only the 6,165-element results. Generally, we manually tune and optimize the hyperparameters of the DeepLPI network and choose the number of blocks empirically in the ResNet-based module.

## Results

### Training and evaluation results

#### Evaluation on BindingDB Dataset

We first report loss and metric progression during training on the training dataset from BindingDB database in **Figure 3**. The model training is stopped after 40 epochs once the validation loss stopped decreasing. Training beyond this point would lead to apparent overfitting marked by increase of the validation loss. We calculated the AUROC metric values during the model training, which achieved 0.95 and 0.89 for the training and validation, respectively.

**Figure 3.**
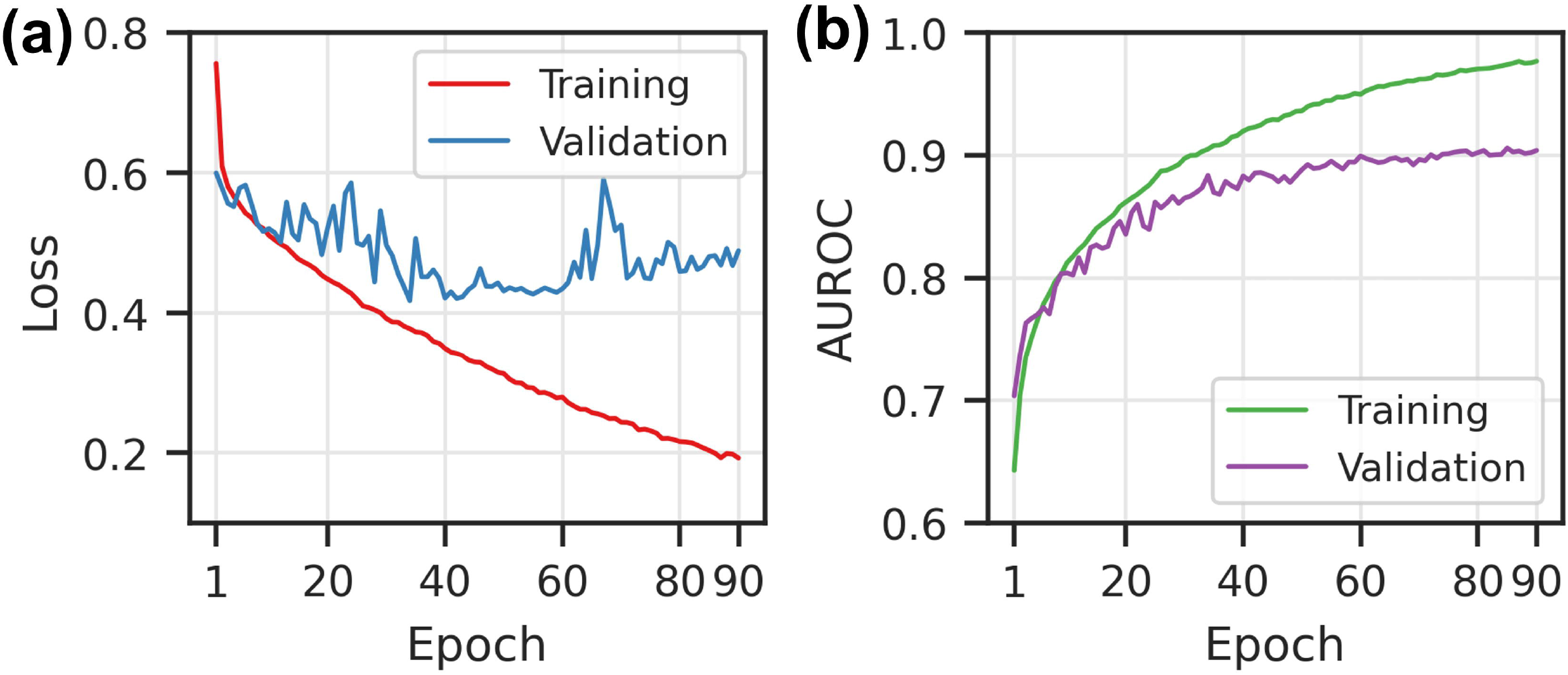
The loss and AUROC score during the DeepLPI training on the BindingDB Kd dataset. **(a)** Loss scores for training and validation. **(b)** AUROC scores for training and validation

We applied the trained model on the internal independent testing set, achieving AUROC measured of 0.893. We used Youden’s J statistic to determine the optimal classification threshold instead of using default value of 0.5 (**Fig. 4a)**. The optimal threshold was used for later calculations of confusion matrix metrics (**Fig. 4b-e**) including the three testing sets “Molecule unseen” (**Fig. 4c**), “Protein unseen” (**Fig. 4d**) and “None seen” (**Fig. 4e**) where the drugs and/or the proteins are of different types (“unseen”) from those in the training set.

**Figure 4.**
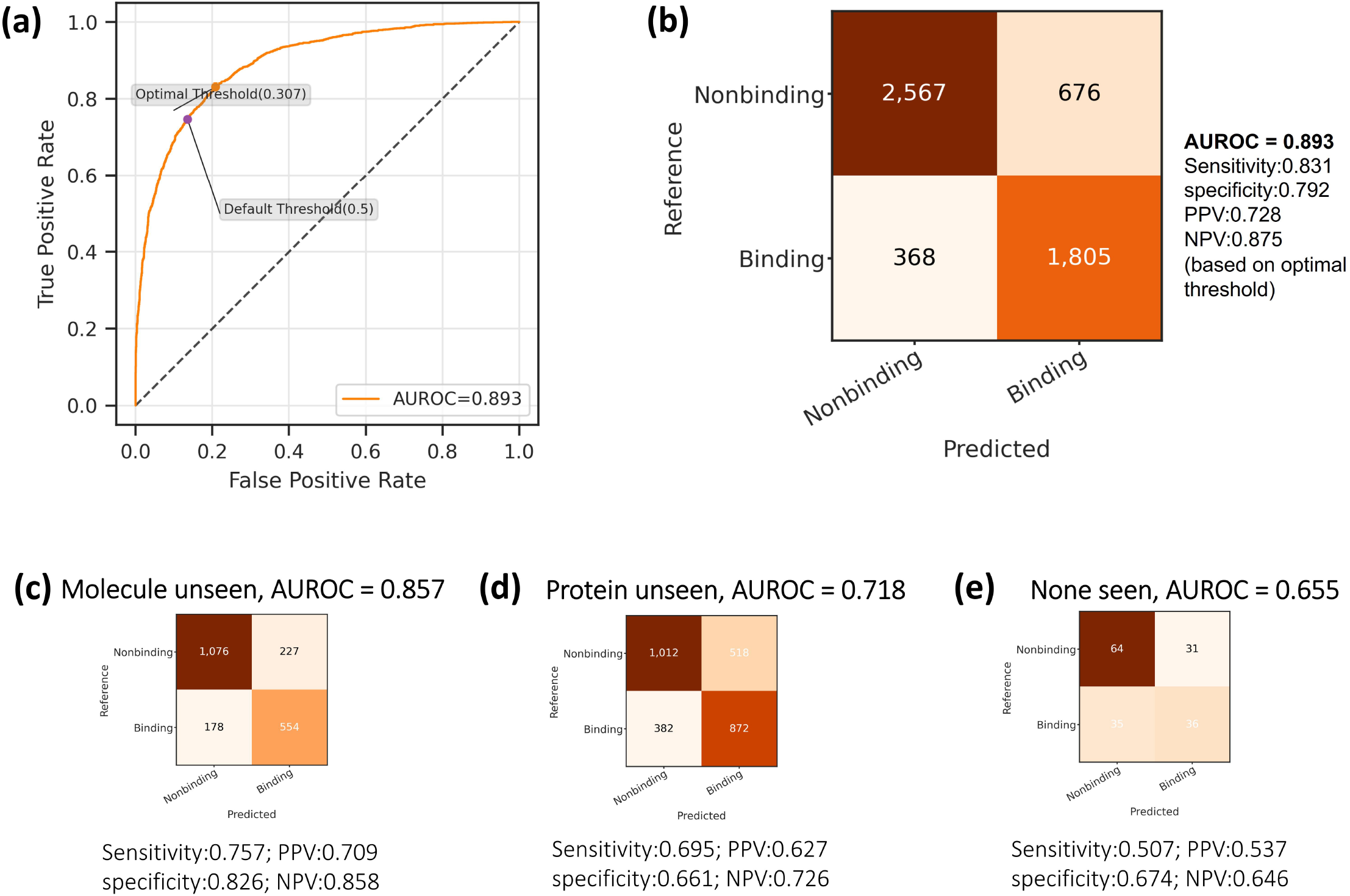
The prediction performance of the final DeepLPI model on BindingDB dataset. **(a)** The ROC curve and the determined optimal threshold. **(b)** Confusion matrix based on the optimal threshold. **(c)** – **(e)** Confusion matrix and performance metrics on the three “unseen” drug/protein testsets: **(c)** Molecule unseen **(d)** Protein unseen and **(e)** None seen.

We observed that the DeepLPI model obtained the highest AUROC of 0.857 when the training set had partial knowledge of the testing set. In addition, AUROC can reach 0.718 when the testing sets contains interaction pairs of molecules (**Fig. 4c**) or protein (**Fig. 4d**) that were “unseen” in the training set. Furthermore, AUROC can achieve 0.655 (**Fig. 4e**) when the training set has no knowledge of the interaction pairs of molecule and protein that exist in the testing set.

In **Table 1**, we compare the performance metrics of our model with the recently published DeepCDA [16] model and the popular baseline model DeepDTA [15] on the independent testing set from the BindingDB data. The results demonstrated that DeepLPI scored higher AUROC by 0.011 and 0.004 than DeepCDA and DeepDTA, respectively. Given that all models’ AUROC are close to 0.9, it can be concluded that DeepLPI is able to predict on BindingDB dataset with very high accuracy. DeepLPI reached sensitivity of 0.832 and NPV of 0.875, which were higher than both DeepCDA and DeepDTA. The specificity and PPV are close to DeepCDA but lower than DeepDTA, due to higher false positive rates indicating the DeepLPI and DeepCDA tend to “over-bind” drug-target pairs. This might result from the more complex network structure with LSTM methods, which invites further investigations.

**Table 1.** Comparing Performance of DeepLPI, DeepCDA and DeepDTA on the internal independent testing set from the BindingDB data.

| **BindingDB** | **AUROC** | **Sensitivity** | **Specificity** | **PPV** | **NPV** |
| --- | --- | --- | --- | --- | --- |
| This work | **0.893** | **0.831** | 0.792 | 0.728 | **0.875** |
| DeepCDA | 0.882 | 0.792 | 0.804 | 0.730 | 0.852 |
| DeepDTA | 0.889 | 0.772 | **0.862** | **0.790** | 0.849 |
| **Unseen Testsets Combined** |  |  |  |  |  |
| This work | **0.790** | **0.684** | 0.773 | **0.671** | **0.783** |
| DeepCDA | 0.448 | 0.000 | **1.0** | Nan | 0.596 |

Overall prediction results on the combined “unseen” testing sets showed DeepLPI with AUROC of 0.790 is 76% better than the DeepCDA model. The DeepCDA with AUROC of 0.448 is non-predictive given that random prediction would lead to AUROC of 0.5, indicating DeepLPI has greater generalizability than the two models when applied on different kinds of drugs and proteins outside of the training domain.

#### Evaluation on Davis dataset

Model training on Davis dataset is stopped after 40 epochs to avoid overfitting when the validation loss stopped decreasing. AUROC values during the model training achieved 0.99 and 0.91 on the training and validation sets, respectively (**Fig. 5**).

**Figure 5.**
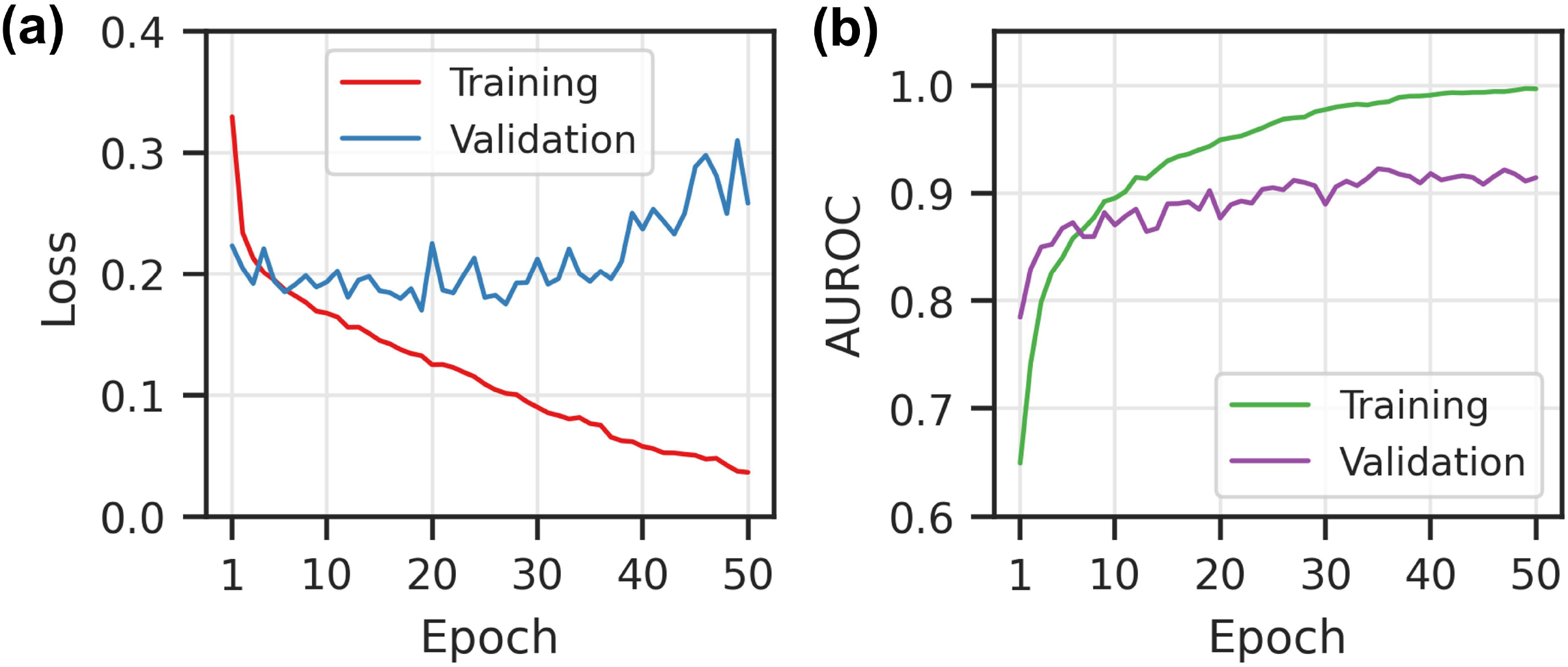
The loss and AUROC score during the DeepLPI training on Davis dataset **(a)** Loss scores for training and validation. **(b)** AUROC scores for training and validation

Evaluating the trained model on the Davis independent testing set, we obtained AUROC of 0.925. We also used Youden’s J statistic to determine the optimal classification threshold (**Figure 6a)**, which was used for later calculation of confusion matrix metrics (**Figure 6b-e**) including “Molecule unseen” (**Fig. 6c**), “Protein unseen” (**Fig. 6d**) and “None seen” (**Fig. 6e**) testing sets. In these “unseen” testing set, DeepLPI model achieved higher accuracy in “Protein unseen” (**Fig. 6d**) testing set with AUROC of 0.812, compared to in the “None seen” (**Fig. 6e**) testing set with AUROC 0.692, and the lowest accuracy was in “Drug unseen” (**Fig. 6c**) testing set with AUROC 0.618. “Drug unseen” was expected to result in higher AUROC because of the partial protein knowledge it includes, and the lower AUROC result might arise from the specific drug molecules collection because Davis dataset did not contain sufficient drug molecules. Larger dataset with more drug molecules could lead to better performance as in the BindingDB case.

**Figure 6.**
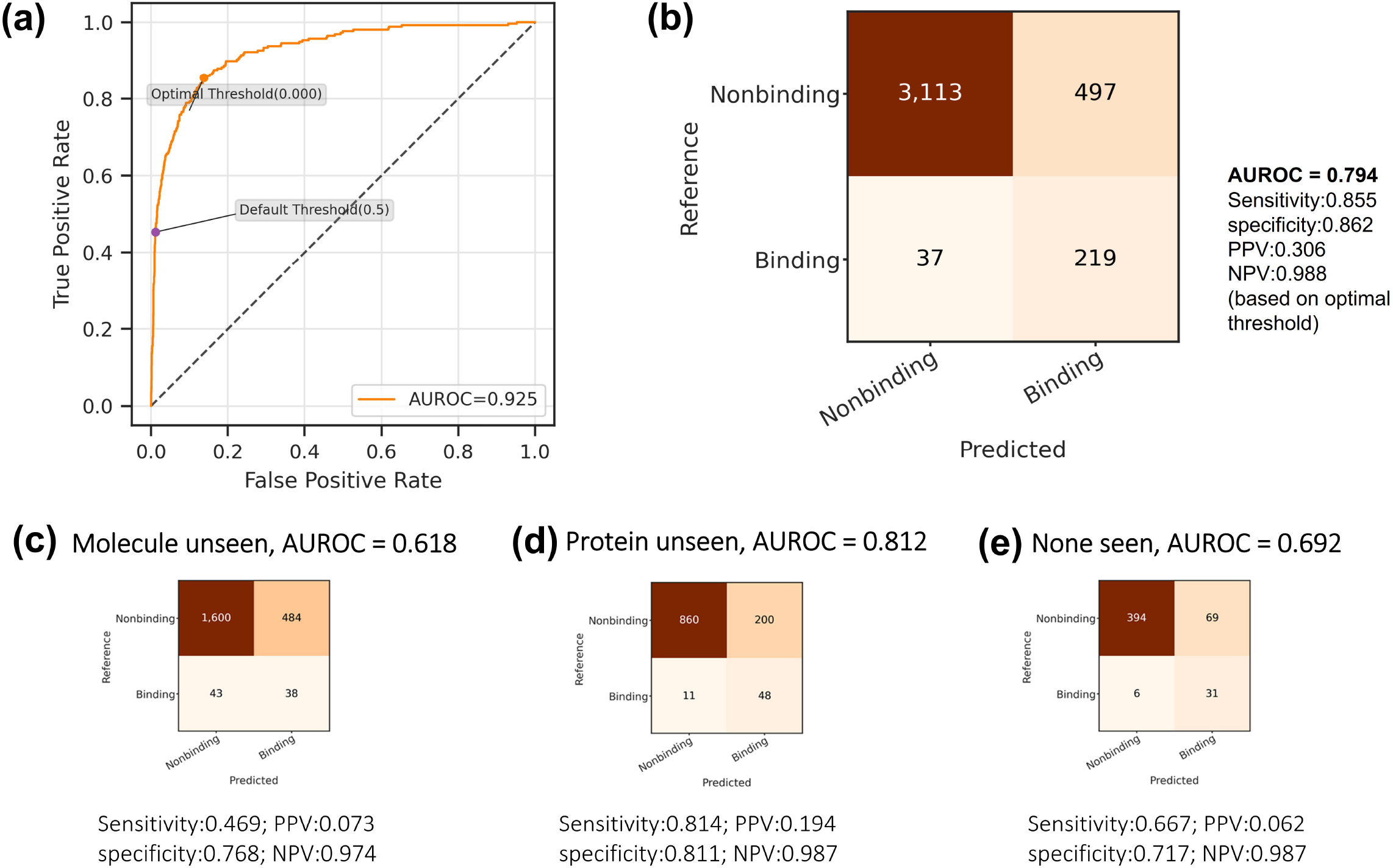
The prediction performance of the final DeepLPI model Davis dataset. **(a)** The ROC curve and the determined optimal threshold. **(b)** Confusion matrix based on the optimal threshold. **(c) – (e)** Confusion matrix and performance metrics on the three unseen drug/protein testsets: **(c)** Molecule unseen **(d)** Protein unseen and **(e)** None seen.

In **Table 2**, we compared the performance metrics of our model with DeepCDA and DeepDTA on the independent testing set from the Davis dataset. DeepLPI with AUROC of 0.925 scored higher values by 0.013 and 0.006 than DeepCDA and DeepDTA, respectively. Given that all models’ AUROC are above 0.9, it can be concluded that DeepLPI is able to predict on Davis dataset with very high accuracy. DeepLPI reached sensitivity of 0.855, specificity of 0.862, NPV of 0.988 and PPV of 0.306, which were higher than or comparable to both DeepCDA and DeepDTA. The low value of PPV may be due to the unbalanced distribution of Davis dataset with less than 10% strong binding entries, and therefore leading to a much smaller true positive rate compared with the false positive. We argue that the result does not invalidate the three models but put into question the suitability of Davis dataset as a benchmark. Overall prediction results on the combined “unseen” testing sets show DeepLPI with AUROC of 0.791 is higher than the DeepCDA model, indicating DeepLPI has better generalizability.

**Table 2.** Comparing Performance of DeepLPI, DeepCDA and DeepDTA on the internal independent testing set from the BindingDB data

|  | **AUROC** | **Sensitivity** | **Specificity** | **PPV** | **NPV** |
| --- | --- | --- | --- | --- | --- |
| DeepLPI | **0.925** | 0.855 | 0.862 | 0.306 | **0.988** |
| DeepCDA | 0.912 | 0.766 | **0.896** | **0.342** | 0.982 |
| DeepDTA | 0.909 | **0.865** | 0.795 | 0.221 | **0.989** |
| **Unseen Testsets Combined** |  |  |  |  |  |
| This work | **0.791** | **0.661** | 0.789 | 0.132 | **0.980** |
| DeepCDA | 0.741 | 0.511 | **0.813** | **0.495** | 0.823 |

#### Evaluation on COVID-19

We further applied the trained models using BindingDB dataset directly on the COVID-19 dataset without any fine-tuning. DeepLPI outperforms DeepCDA with an AUROC of 0.610 (**Table 3**). The high prediction performance on COVID-19 dataset suggests that our DeepLPI may be the potential method to find effective drugs for SARS-CoV-2. The low PPV and specificity of DeepLPI arise from large false positive rates and are indicating the potential upgrades in our future works.

**Table 3.** Comparison of DeepLPI and DeepCDA on transferring BindingDB trained model to COVID-19.

|  | **AUROC** | **Sensitivity** | **Specificity** | **PPV** | **NPV** |
| --- | --- | --- | --- | --- | --- |
| DeepLPI | **0.610** | 0.538 | 0.576 | **0.110** | **0.928** |
| DeepCDA | 0.400 | 0.000 | **1.000** | nan | 0.911 |

## Discussion

In our work, we successfully build DeepLPI model to predict DTI in classification tasks using 1D sequence data from protein and drug molecules. We first utilize the pre-trained embedding methods called Mol2Vec and ProSE to encode the raw drug molecular SMILES strings and target protein sequences respectively into dense vector representations. Then, we feed the encoded dense vector representations separately into head modules and ResNet-based modules to extract features, where these modules are based on 1D CNN. The extracted feature vectors are concatenated and fed into the biLSTM network, further followed by the MLP module to finally predict binary active or inactive based on *K_d_* affinity labeled data. We used three datasets of BindingDB, Davis and COVID-19 to evaluate our DeepLPI model, and the results demonstrate that our model has a high performance on the prediction.

Unlike the methods to pre-define features that are heavily relied on domain knowledge or to represent sequences simply using sparse encoding approach, our DeepLPI applied pretrained embedding models of Mol2Vec[17] and ProSE[18] to encode the raw drug SMILES string and target protein sequences, respectively. These semantic context embedding models are trained using a huge dataset to represent sequence data in the form of dense vectors, with consideration of structure information of molecule and target proteins to ensure that they are highly informative and efficient for feature embeddings. It is admired that there exists a variety of embedding methods to encode drug compounds and protein sequences, we picked Mol2Vec and ProSE in our DeepLPI due to our empirical experience. We use 1D CNN in our DeepLPI model to retain the sequential correlation. We adopt a ResNet-based module in the DeepLPI. Traditional feed-forward CNN may lose useful information as the design grows deeper. Nevertheless, ResNet-based CNN can mitigate this drawback by developing a “shortcut connection” for the network. Consequently, data inputted into the ResNet-based CNN module can be added with the residual of the network to alleviate the loss of information. The biLSTM is employed in the DeepLPI model, which can capture long-term dependencies of the sequence, equally encode input sequence once from beginning to end and once from end to beginning. Compared to the classical LSTM, the biLSTM enables the use of the two hidden states in each LSTM memory block to preserve information from both past and future.

In our experiments, we notice that the performance of DeepLPI is not uniform on different proteins: there might exist some common biological features of those proteins such as the sequences or the spatial structures. Detailed analysis of the shared features of the proteins requires a deeper understanding of the protein-drug interaction and can potentially explain the reason that the model behaves well on some of the proteins. Such analysis would be useful to improve the model when we generalize the results later.

The DeepLPI model may help in speeding up the COVID-19 drug research. As of today, the pandemic is not showing any sign of slowing down and people are still searching for an effective and safe cure for COVID-19 patients. The current widely-used combination treatment with hydroxychloroquine and azithromycin has not been proven to be satisfactory, and there are some research efforts in using computational, especially deep neural network, techniques for searching the effective repurposed drugs. Our model can be useful in speeding up the drug search and potentially increase the success rate because the training data fed into the model is not limited to the protein structural information.

Even though we have successfully built a model that can predict active/inactive interaction with high accuracy, the model still suffers from some limitations. There is still room for improvement regarding the prediction accuracy, especially when the model is applied on external datasets. From a broader perspective, the study of repurposing drugs should not be limited only to the binding affinities. Researchers should also pay attention to the possibility of potential adverse effects of using the repurposed drug. This can be a result of new interactions between the drug and the proposed disease target, or because the drug is administered to a new group of population. Sometimes the repurposed drug could have interactions with traditional drugs on the new disease, and adverse effects might also arise from such unexpected interactions. Deep learning methods could also be used in studying on these aspects for better safety.

## Acknowledgements

B. W. would like to thank Dr. Yingce Xia for insightful discussions on the embedding algorithms and the testing methods for generalizability. B. W. would like to thank Dr. Yong Bai for his advice in deep learning algorithms.

## Data Availability

The datasets including drug information and protein information analyzed during the current study are publicly available online through the following links:

BindingDB data (http://www.bindingdb.org/bind/chemsearch/marvin/SDFdownload.isp?all_download=yes),

Davis data (http://staff.cs.utu.fi/~aatapa/data/DrugTarget/), and COVID-19 data (https://www.diamond.ac.uk/covid-19/for-scientists/Main-protease-structure-and-XChem.html)

The code of the LPI model and the training and testing script notebooks are available open source through the following Github link (https://github.com/David-BominWei/DeepLPI).

## Author Contributions

B. W. conceived the idea. All authors contributed to the proposal of the algorithm and selection of database and baseline methods. B. W. and X. G. contributed to the choice of metrics. B. W. and Y. Z. contributed to the construction of testing methods. B. W. wrote the manuscript and all authors contributed to the manuscript revisions.

---

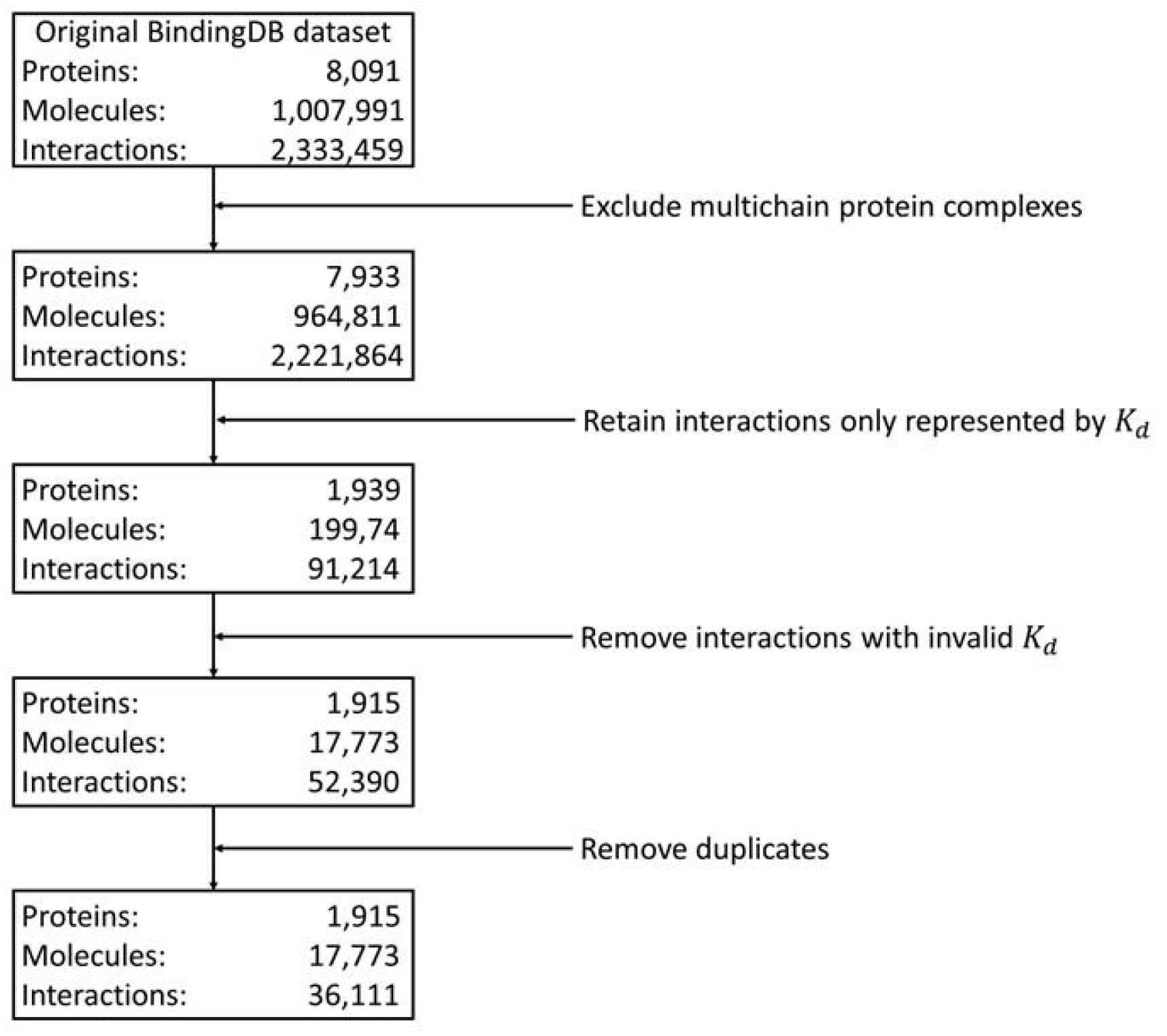

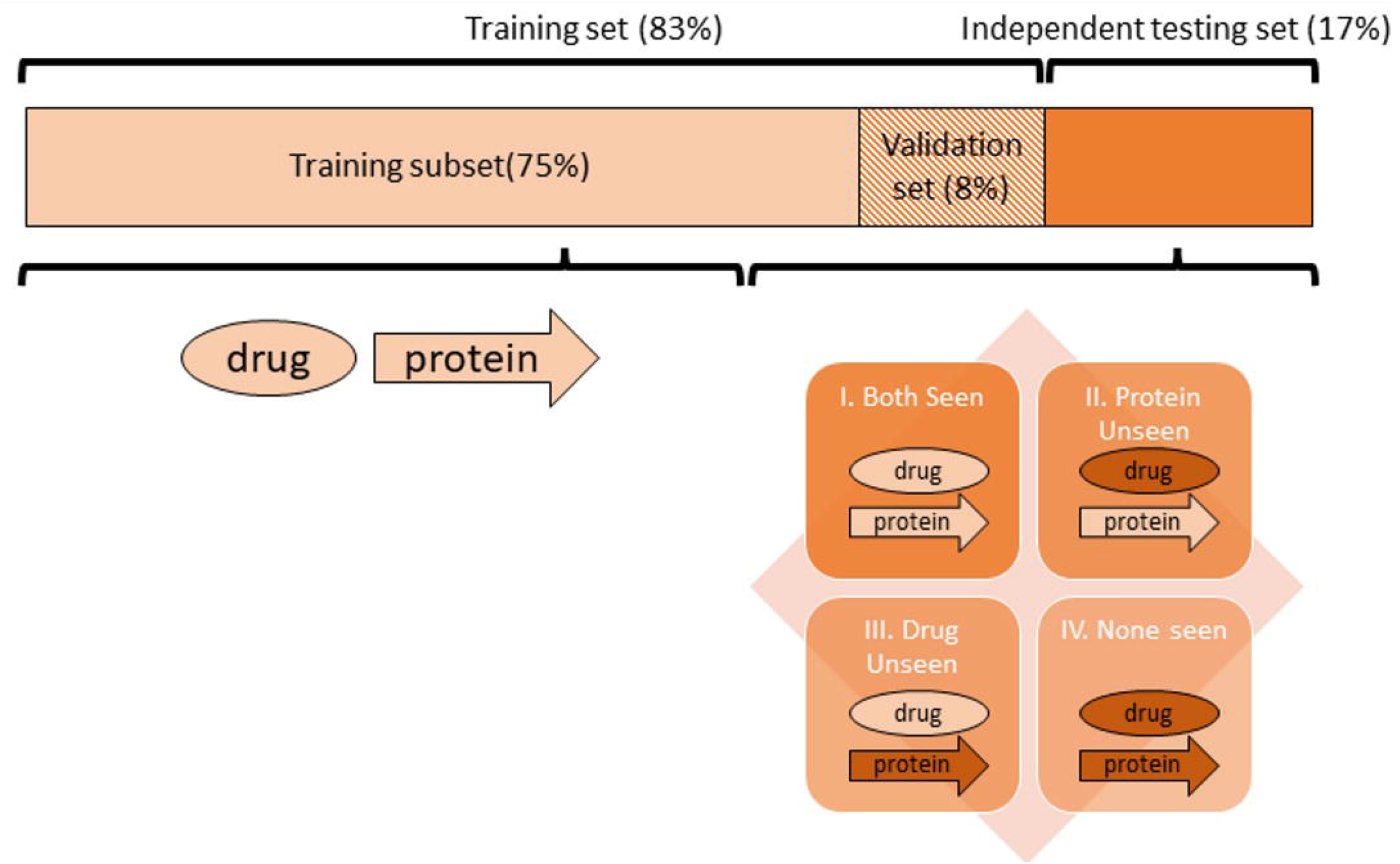

