## Supplementary table and figure for "DeepLPI: a novel deep learning-based model for protein-ligand interaction prediction for drug repurposing"

1. Parameter settings for DeepLPI model training

**Table S1** The parameter settings for the DeepLPI

|  |  | Drug compounds | Target proteins |
| --- | --- | --- | --- |
| Modules | Parameters | Value | Value |
| Head Module | Number of kernels | 32 | 32 |
|  | Kernel size | 7 | 7 |
|  | Stride | 2 | 2 |
|  | Padding | 3 | 3 |
| ResNet-based CNN module | Number of kernels | [32,32], [16,16], [16, 16] | [32,32], [16,16], [16, 16] |
|  | Kernel size | [3,3], [3,3], [3,3] | [3,3], [3,3], [3,3] |
|  | Stride | 1 | 1 |
|  | Padding | 1 | 1 |
| Max Pooling 1D | Kernel size | 2 | 2 |
|  | Stride | 2 | 2 |
| Average Pooling 1D | Kernel size | 5 | |
|  | Stride | 3 | |
| biLSTM module | Input size | 538 | |
|  | Hidden size | 64 | |
|  | Number of layers | 2 | |
|  | Bidirectional | True | |
| MLP module | Number of neurons | [2048,512,32] | |
| Common parameter setting for all modules | Dropout | 0.3 | |
|  | Weight initialization | Kaiming | |
|  | Optimizer | Adam | |
|  | Batch size | 256 | |
|  | Learning rate (LR) | 0.001 | |
|  | Weight for L2-norm ($\alpha$) | 0.0001 | |
|  | LR decay rate | 0.8 | |

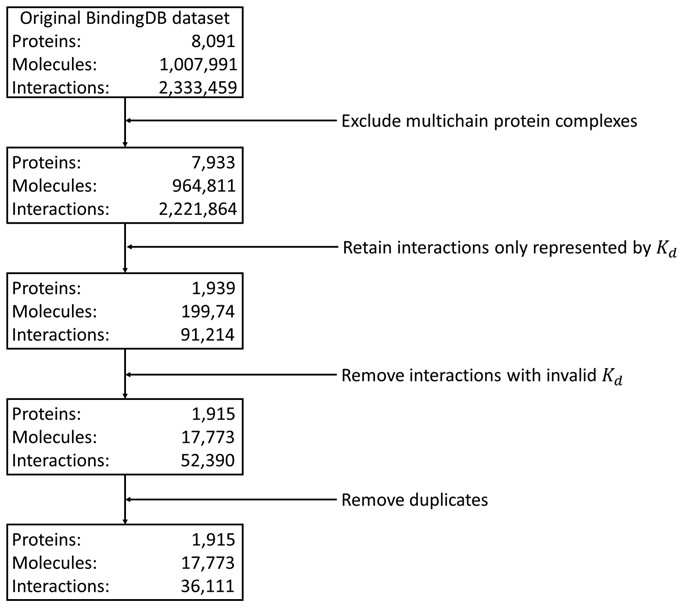
 **Figure S1** Preprocessing of BindingDB dataset. Data exclusion criteria to compile BindingDB dataset.

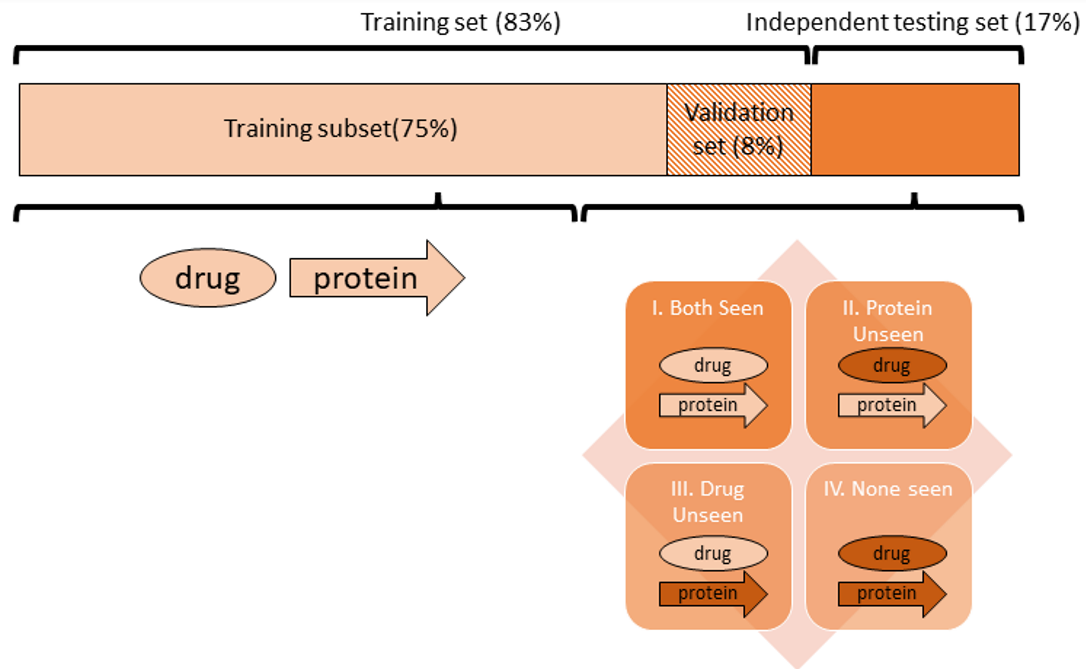
**Figure S2** Construction of the unseen testing sets. Split the whole dataset into a training set ($83\%$ of whole interactions) and an independent testing set ($17\%$) for training and evaluating the model, respectively. The training set is further divided into the training subset ($75\%$) and the validation set ($8\%$). The independent testset was further splitted into for parts. Part I: the drug or protein information is separately included in the training set but not their pairs. Part II: the drug information included in the training set but not the protein. Part III: the protein information is included in the training set but not the drug information. Part IV: neither of drug or protein information is included in the training set.
